# Specialized and experience-dependent neural dynamics in the medial amygdala and bed nucleus of stria terminalis upon social and sexual olfactory and tactile cues

**DOI:** 10.64898/2026.10.23.761408

**Authors:** Patty T. Huijgens, Roy Heijkoop, Eelke M.S. Snoeren

**Affiliations:** Department of Psychology, UiT the Arctic University of Norway, Tromsø, Norway

**Keywords:** medial amygdala, bed nucleus of stria terminal, sexual behavior, social behavior, ejaculation, rats, males, neural patterns

## Abstract

The medial amygdala (MeApd) and bed nucleus of the stria terminalis (BNST) are central components of the social behavior network. However, their real-time activity during social and sexual interactions, and their respective contributions to processing olfactory and tactile cues, remain unclear. Here, we used simultaneous dual-site fiber photometry to record GCaMP6s signals from the MeApd and BNST in freely behaving male rats during social interaction, copulation, sexual exhaustion, and vaginal occlusion tests. Introduction of a conspecific increased activity in both regions, but the response profiles differed by social context and experience. The MeApd responded robustly to male and female conspecifics, with larger responses to males, whereas BNST activity increased reliably upon female introduction only after sexual experience. During olfactory investigation of a male conspecific, MeApd activity differentiated sniffing of anterior and posterior body regions, while BNST activity showed no comparable spatial selectivity. Neither region showed robust event-aligned responses during sniffing of receptive females. Tactile stimulation during copulation elicited strong calcium responses in both regions. Mounts, intromissions, and ejaculations each triggered significant activity, with ejaculation producing the largest responses. MeApd responses differentiated ejaculation from mounting in sexually naïve animals, whereas distinctions in BNST responses emerged primarily after sexual experience. Moreover, responses to intromissions and ejaculations persisted during the final ejaculatory series preceding sexual exhaustion, while responses to mounts were eliminated when vaginal penetration was prevented. Together, these findings identify specialized yet parallel activity dynamics within the MeApd–BNST network. MeApd activity preferentially tracks socially relevant olfactory investigation and the intensity of copulatory sensory feedback, whereas BNST recruitment during sexual behavior is more strongly shaped by sexual experience.

## Introduction

Many animals, including humans and rodents, are intrinsically social organisms that rely on complex group dynamics for survival, reproduction, and overall well-being^1–3^. Social interactions require the brain to identify conspecifics, evaluate their biological relevance, and select context-appropriate behavioral responses. These interactions involve complex sequences of integration of sensory cues leading to approach and interaction, which in sexual behavior can lead to copulation and ejaculation. The behaviors rely on the integration of sensory, hormonal, and contextual information, mediated by distinct yet interconnected neural circuits ^4–7^.

Social and sexual behavior in rodents begins with investigation of conspecifics, often involving sniffing of pheromone-rich facial and anogenital regions ^8^. Chemosensory information is conveyed through the main and accessory olfactory systems to the medial amygdala (MeA) and bed nucleus of the stria terminalis (BNST), which in turn communicate with hypothalamic and mesolimbic systems involved in reproductive and social behavior ^6,9–15^. Although olfactory cues are major determinants of social and sexual motivation in rodents^16^, social investigation also provides tactical stimulation through physical contact. Although the specific role of tactile cues in rodent social and sexual investigation remains poorly understood, tactile input serves fundamental homeostatic functions, and disrupted tactile feedback impairs social motivation during isolation^17^. Moreover, sexual touch can powerfully influence sexual development^18^, while sensory feedback from genital stimulation is linked to hedonic sensation and sexual performance^19–21^. During copulation, genital tactile feedback becomes especially prominent and may differ in intensity and sensory consequences across mounting, intromission, and ejaculation. The pathways through which genital and other tactile signals influence sociosexual behavior remain incompletely understood. Tactile information from cutaneous low-threshold mechanoreceptors ascends via the spinal cord and thalamus to the somatosensory cortex^22^, where socially relevant signals are integrated within hypothalamic circuits centered in the medial preoptic area, which communicates extensively with the ventral tegmental area, nucleus accumbens, MeA, and BNST ^12,13^.

Beyond routing olfactory and tactile cues, the MeA and BNST are directly involved across phases of social and sexual behavior. Immediate-early genes in these regions are induced by anogenital investigation, exposure to odors from receptive females, and copulation^23–27^. Neural activity recordings in rats and mice show robust MeA and BNST activation upon introduction of a receptive female, with further spikes following mounts, intromissions, and ejaculation^28,29^. Although the MeA and BNST clearly respond during copulation, lesions or chemogenetic inhibition do not abolish copulation or ejaculation. In the MeA, they typically prolong ejaculation latency and increase the number of mounts and intromissions preceding ejaculation ^30–36^, whereas in the BNST they impair sexual performance by prolonging ejaculation latency, increasing inter-intromission intervals, and extending post-ejaculatory refractory periods ^37–39^. This suggests that the BNST coordinates the temporal dynamics of copulatory behavior, while the MeA integrates olfactory/pheromonal cues and genital somatosensory feedback to modulate ejaculatory responses^25,36,40^.

Despite this evidence, the real-time dynamics of MeA and BNST activity across social investigation and discrete copulatory events remain poorly understood in rats. In particular, it is unclear whether these interconnected regions show distinct activity patterns during social versus sexual investigation, whether their responses differ across copulatory events, and how sexual experience modifies their recruitment. Rats provide an advantageous model for addressing these questions, because they have a highly social behavioral ecology and their copulatory repertoire is well characterized and includes identifiable mounts, intromissions, ejaculations and progressive sexual satiety after repeated ejaculations^41,42^. In addition, while most forms of social behavior, such as infant-adult bonding and peer-to-peer interactions, emerge immediately after birth, sexual behavior represents a more specialized form of social interaction that develops only after sexual maturation^43,44^. This developmental distinction provides a unique opportunity to study transition from sexually naïve to experienced states and compare sexual behavior with other forms of social interaction.

Here, we used simultaneous dual-site fiber photometry to monitor bulk calcium dynamics in the posteromedial nucleus of the MeA (MeApd) and BNST of male rats during social interaction, sexual incentive motivation, copulation, sexual exhaustion, and copulation with vaginal occlusion. We tested whether MeApd and BNST activity differed during social and sexual investigation, whether neural activity associated with copulatory events varied across mounts, intromissions, and ejaculations, and whether these activity patterns were modified by sexual experience. By comparing two interconnected extended-amygdala nodes in the same animals, this study provides a framework for examining how social context, copulatory sensory feedback, and experience shape neural activity during social behavior.

## Material and Methods

### Rats

Fourteen experimental male Wistar rats were implanted for dual-site fiber photometry. Seven animals with histologically verified GCaMP6s expression and fiber placements in both the MeApd and BNST were included in the final analyses. Four intact males and eight ovariectomized females served as social and sexual stimulus animals, respectively.

Rats were obtained from Janvier Labs (France) at a body weight of 200–240 g. Animals were pair-housed in Macrolon IV® cages under a reversed 12-h light/dark cycle, with lights on from 23:00 to 11:00 h. Food (low phytoestrogen maintenance diet, #V1554, Ssniff, Germany ) and water were available ad libitum. Temperature and humidity were maintained at 21 ± 1 °C and 55 ± 10%, respectively. All procedures complied with European Union Directive 2010/63/EU and were approved by the Norwegian Food Safety Authority (FOTS #22687) on the 28^th^ of April 2020.

Animals were excluded when viral expression or fiber placement was outside the intended target in either the MeApd or BNST (Fig. S1).

### Surgical procedures

#### Brain surgery

All experimental males underwent stereotaxic surgery for viral-vector infusion and optic fiber implantation. Rats were anesthetized with isoflurane and placed in a stereotaxic apparatus (Stoelting Europe, Ireland). Buprenorphine (0.05 mg/kg, s.c.) and meloxicam (2 mg/kg , s.c.) were administered for analgesia. The scalp was shaved, disinfected with povidone-iodine and locally anaesthetized with bupivacaine (0.15 ml, s.c.). Core body temperature, oxygen saturation and heart rate were monitored throughout the surgery.

AAV5-hSyn-GCaMP6s (pAAV5.Syn.GCaMP6s.WPRE.SV40, plasmid #100843 from Addgene) was infused into the right MeApd and BNST. Two additional animals received AAV5-hSyn-eGFP (pAAV5.hSyn.eGFP.WPRE.bGH, plasmid # 105539 from Addgene) as fluorophore controls. In each target region, 500 nl virus was infused at 150 nl/min, followed by a 10-min diffusion period before needle withdrawal. This was done using a 10 µl Hamilton syringe with a 30G blunt needle mounted in Hamilton injectors (Stoelting) on the stereotaxic apparatus. Optical fibers (diameter 400 µm, NA 0.66, Doric Lenses, USA) were implanted at the same coordinates used for viral infusion (fiber length: BNST: 8.0 mm and MeA: 10.0 mm) and secured with anchor screws (1 x 3 mm, Agnthos, Sweden), Super-Bond adhesive (CCB Super bond, Technomedics, Norway), and dental acrylic (Meliodent, Kulzer GmbH, Germany).

For the MeApd, the starting point of insertion of the Hamilton syringe needle and the optic fiber was AP −3.1 mm, ML +2.60 mm relative to bregma, with an 8° coronal angle with the fiber tip pointing towards lateral. Insertion depth was 9.20 mm, aiming for the coordinates AP -3.1, ML +4.1, DV -8.1. For the BNST, the insertion location was AP +1.35 mm, ML +1.25 mm with a 16° sagittal angle with the fiber tip pointing towards posterior. Insertion depth was 6.75 mm, aiming at AP -0.8, ML +1.25, DV -6.1. Fiber lengths were 10.0 mm for MeApd and 8.0 mm for BNST. Rats were post-operatively treated with analgesia consisting of 0.05 mg/kg buprenorphine and 2 mg/kg meloxicam after 24 and 48 hours, and immediately pair-housed after surgery. A period of four weeks was used for virus expression before the behavioral experiments started.

#### Ovariectomy

Two weeks before the experiments, all stimulus females were ovariectomized under isoflurane anesthesia as previously described^45^. Briefly, a medial dorsal incision of the skin of about 1 cm, and small incisions in the muscle layer on each side, were made. The ovaries were located and extirpated, and the muscle layer sutured. A silastic capsule containing β-estradiol was placed subcutaneously. The skin was closed with a wound clip. Females were allowed at least one week of recovery before use in behavioral tests. Analgesic treatment consisted of 0.05 mg/kg buprenorphine and 2 mg/kg meloxicam pre-operative and post-operative after 24 and 48 hours.

### Hormonal priming of stimulus females

The silastic capsules (medical grade Silastic tubing, 0.0625 in. inner diameter, 0.125 in. outer diameter; Degania Silicone, Degania Bet, Israel) for females were 5 mm long and contained 10% 17β-estradiol (E8875, Sigma, St. Louis, USA) in cholesterol (C3292, Sigma, St. Louis, USA). The silastic tubing was closed off by inserting pieces of toothpick into both ends and sealed with medical grade adhesive silicone (A-100; Factor II Incorporated, Arizona, USA). Progesterone (P0130, Sigma, St. Louis, USA) was dissolved in peanut oil (Apotekproduksjon, Oslo, Norway) at a concentration of 5 mg/mL and injected subcutaneously with 0.2 ml of 1 mg 4 hours before behavioral testing in a sexual context.

### Behavioral assessment

#### Sexual incentive motivation

The sexual incentive motivation (SIM) test (see also:^46^) apparatus is a rectangular arena (100 × 50 × 45 cm) with rounded corners with incentive cages attached with a wired mesh (25 × 25 cm) to the arena in opposite corners of the long sides. All rats were first habituated to the test arena for 10 minutes on three different days the week before the experiment started with no incentive stimuli present. During normal testing, a social stimulus (intact male rat) was placed in one of the stimulus cages and a sexual stimulus (receptive female rat; ovariectomized, implanted with a 10% 17β-estradiol silastic capsule, and injected with 1 mg progesterone 4 h before use in the test (standard procedure in our laboratory^16,36,47^) was placed in the other stimulus cage. The position of the stimulus cages and the stimulus rats were randomly changed throughout each experimental session. The experiment starts with the placement of the subject male in the middle of the arena. For 10 minutes, the x and y coordinates of both the center and the nose of the subject were determined by video-tracking with Ethovision software (Noldus, Wageningen, the Netherlands). With the use of a customized Python 3.13 script (available upon request), both the entering and exiting of the two virtual stimulus zones (30 × 21 cm) in front of each stimulus cage (based on the center of the subject) and the approach of the stimuli next to the wired mesh (based on the nose coordinates) were marked. From this, the time spent in each incentive zone and the preference score was calculated by dividing the time spent in the female incentive zone by the total time spent in incentive zones. To control for locomotor behavior artifacts, the distance moved and mean velocity were also measured during the test. The tests were conducted in dimly lit (ca 5 lx) conditions. Between tests, feces and urine were removed from the arena, and the SIM arena was cleaned with diluted acetic acid between experimental days.

#### Copulation

The copulation (COP) test was conducted in rectangular boxes (40 × 60 × 40 cm) with a Plexiglas front, in a dimly lit (ca 5 lx) room. The bedding always contained some odors of receptive females from previous tests. At the time of testing, the male subject was allowed 5 minutes of habituation to the copulation box, after which a receptive female (injected with 1 mg progesterone 4 hours before use) was introduced, which started the 30 minute test. All copulation tests were recorded on camera and behavior was later assessed from video. Behavioral assessment consisted of scoring behavioral events in Observer 17XT software (Noldus, Wageningen, the Netherlands). For the entire 30 min, the copulatory behaviors mount (pelvic thrusting while being mounted on the female), intromission (mounting the female with pelvic thrusting and penile insertion into the vagina; characterized by a more vigorous dismount) and ejaculation (characterized by a longer intromission and the female moving away from the male) were scored. In addition, other behaviors such as genital grooming (autogrooming of the genital region), other grooming (autogrooming in other regions than the genitals), chasing (running after the female), sniffing the female (differentiated the anterior (head/face) versus the posterior part of the body (anogenital/hind area)), sniffing the bedding, rearing (lifting two paws up in the air), were also annotated.

#### Social interaction

Social interaction was tested in the same rectangular boxes (40 × 60 × 40 cm) with a Plexiglas front as the COP test, in a dimly lit (ca 5 lx) room. The cages were carefully cleaned before the test to remove any odors of sexually receptive animals, and new clean bedding was added. At the time of testing, the male subject was allowed 5 minutes of habituation to the social interaction box, after which another male was introduced for 10 minutes. All social interaction tests were recorded on camera and behavior was later assessed from video and annotated with video frame precision, as described above in the COP test. The same behaviors as during the copulation test were scored (if they occurred).

#### Sexual exhaustion

In this test, copulation behavior was tested until sexual exhaustion was reached. A regular COP test was used, except that the males were allowed to continue copulating until they didn’t reach another ejaculation within 90 minutes after the last ejaculation. 24 Hours after the sexual exhaustion test, the males were re-exposed to a regular COP test of 30 minutes.

#### Copulation with vaginal occlusion

In the final behavioral test, males copulated with receptive females whose vaginal opening was covered with sterile gauze and medical tape. This manipulation prevented penile penetration while allowing mounting behavior. Mounts and extravaginial intromission-like patterns ^48^were scored and aligned to photometry signals.

### Fiber photometry recordings and analysis

Bulk fluorescence signals from GCaMP6s-expressing cells were recorded using a Tucker-Davis Technologies RZ10x system. Excitation light at 465 nm was used to measure calcium-dependent fluorescence, whereas 405-nm excitation served as an isosbestic reference channel. Signals were modulated at 210 Hz and 330 Hz, respectively, filtered and combined by a fluorescence mini cube (Doric Lenses, USA), transmitted through low-autofluorescence fiber optic patch cable (Doric, MFP_400/430/1100-0.57_5m_FCM-MF2.5_LAF) to implanted fibers (Doric, MFC_400/430-0.66_9mm_MF2.5_FLT), and acquired using Synapse software (Tucker-Davis Technologies, USA). As we tested two brain regions simultaneously within 1 rat, we used two different fiber optic patch cables, each connected to one of the implanted fiber cannulas. LED power was set at a level which resulted in a ∼15 and ∼30 mV amplitude output signal for 405 and 465 nm demodulated channels, respectively. Fluorescence was sampled at 1017 Hz and demodulated by the processor.

Behavioral-test onset, incentive-zone entries, and video recordings were synchronized with the photometry system using TTL pulses generated by the USB-I/O box 12 system (Noldus, The Netherlands) linked to Ethovision (Noldus, The Netherlands). Behavioral events scored in The Observer XT (Noldus, The Netherlands) were aligned to the photometry data using these timestamps.

The 405-nm signal was linearly fitted to the 465-nm signal, and ΔF/F was calculated as: ΔF/F = (465 nm signal − fitted 405 nm signal)/fitted 405 nm signal^49^. Recording segments containing clear mechanical artifacts or signal disruptions (e.g., when the cable detached or hit the wall or door) were removed from the recordings when an interquartile range of more than 4 was found. This method performed best in the Dickey-Fuller and crest factor tests to maintain high quality data ^50^. Signals were baseline-corrected relative to the pre-event period. Area under the curve, peak amplitude, and response onset were calculated from event-aligned traces.

### Brain processing, immunostaining and imaging

At the end of the experiments, rats were injected intraperitoneally with a lethal dose of pentobarbital (100 mg/kg), and transcardially perfused with 0.01 M phosphate buffered saline (PBS; pH 7.4) followed by 4% formaldehyde in 0.01 M PBS. Brains were removed and post-fixed in 4% formaldehyde for 48 hours. Then, the brains were sectioned on a vibratome (Leica VT1200s)(SM2010R, Leica Biosystems, Germany) into 50 µm coronal sections. These sections were stored in cryoprotectant solution (30% sucrose w/v, 30% ethylene glycol v/v in 0.01 M PBS, pH 7.4) until further use.

To assess the expression of GCaMP6s, immunohistochemistry was performed on 1 in every 3 sections throughout and surrounding the MeApd and BNST. These were washed in 0.1M Tris-buffered-saline (TBS), blocked for 30 minutes in 0.5% BSA in TBS, and incubated with polyclonal chicken anti-EYFP (1:100 000), Abcam, cat. ab13970) antibody solution containing 0.1% Triton-X and 0.1% BSA in TBS on an orbital shaker for 24 hours at room temperature + 24 hours at 4 °C. The sections were then washed four times in TBS and subsequently incubated in biotinylated goat anti-chicken (1:400, Abcam, cat. ab6876) antibody solution containing 0.1 % BSA in TBS for 30 min, followed by incubation with avidin-biotin-peroxidase complex (VECTASTAIN ABC-HRP kit, Vector laboratories, cat. PK-6100, dilution: 1 drop A + 1 drop B in 10 mL TBS) solution for 30 min, and 3,3′-diaminobenzidine solution (DAB substrate kit (HRP), Vector laboratories, cat. SK-4100, dilution: 1 drop R1 (buffer solution) + 2 drops R2 (3,3′-diaminobenzidine solution) + 1 drop R3 (hydrogen peroxide solution) in 5 mL water) for 5-10 min, with TBS washes between all steps. The slides were left until dry, after which the sections were dehydrated with 50-99.8% ethanol baths, cleared with xylene, and coverslipped using Entellan mounting medium (Sigma, St. Louis, USA).

Finally, the accurate placement of the fibers and GCaMP6s expression were verified with a microscope (Primo Star, Carl Zeiss, Germany). Only rats with confirmed expression in both the MeApd and BNST (*n* = 7) and fiber tips in these GCaMP6s expressing regions were included in subsequent analyses. We also confirmed that the two control animals with EYFP, but no GCaMP6s expression, had stable signals and are not included in the rest of the data analysis.

### Statistical analysis

Behavioral and fiber photometry data were analyzed with customized Python 3.13 scripts (available upon request). The data obtained from the copulation tests were combined in two categories: sexually naïve and sexually experienced, with the behaviors from the 1^st^ and 2^nd^ copulation test included in naïve, and the behaviors from the 3^rd^ to 6^th^ copulation test added to the experienced group. Linear mixed models were used to account for the hierarchical structure of the dataset. In the sexual incentive motivation test, an interaction effect between factors Test (naïve versus experienced), Epoch (pre versus post), Incentive (either Female zone entering versus Male zone entering or Female sniffing versus Male sniffing) was calculated for the AUC, and an interaction effect between factors Test and Incentive for the other parameters. For the behavioral parameters in the other tests, a mixed model approach with factors Test (naïve, experienced, and if relevant, social) and Epoch (pre versus post) were used. When different behaviors or the effect of brain region was compared, factor Behavior or Region, respectively, was also used in the model. In case of significance, a Bonferroni post-hoc analysis was performed with α set on 0.05. A paired t-test was performed in cases where two outcomes were compared directly (e.g. the amplitude of the neural response in naïve vs. experienced state), or a ONEWAY ANOVA when 3 conditions were compared (e.g. the amplitude of the neural response between naïve, experienced or social conditions).

## Results

Histological analysis confirmed GCaMP6s expression and optical-fiber placements in both the MeApd and BNST in seven animals, which were included in subsequent analyses (Fig. 1A, Fig. S1). Five of the fourteen GCaMP6s-injected animals were excluded because one or both fiber placements or viral-expression sites fell outside the intended target regions. The two eGFP control animals showed stable fluorescence signals but were not included in the GCaMP6s analyses.

**Fig. 1.**
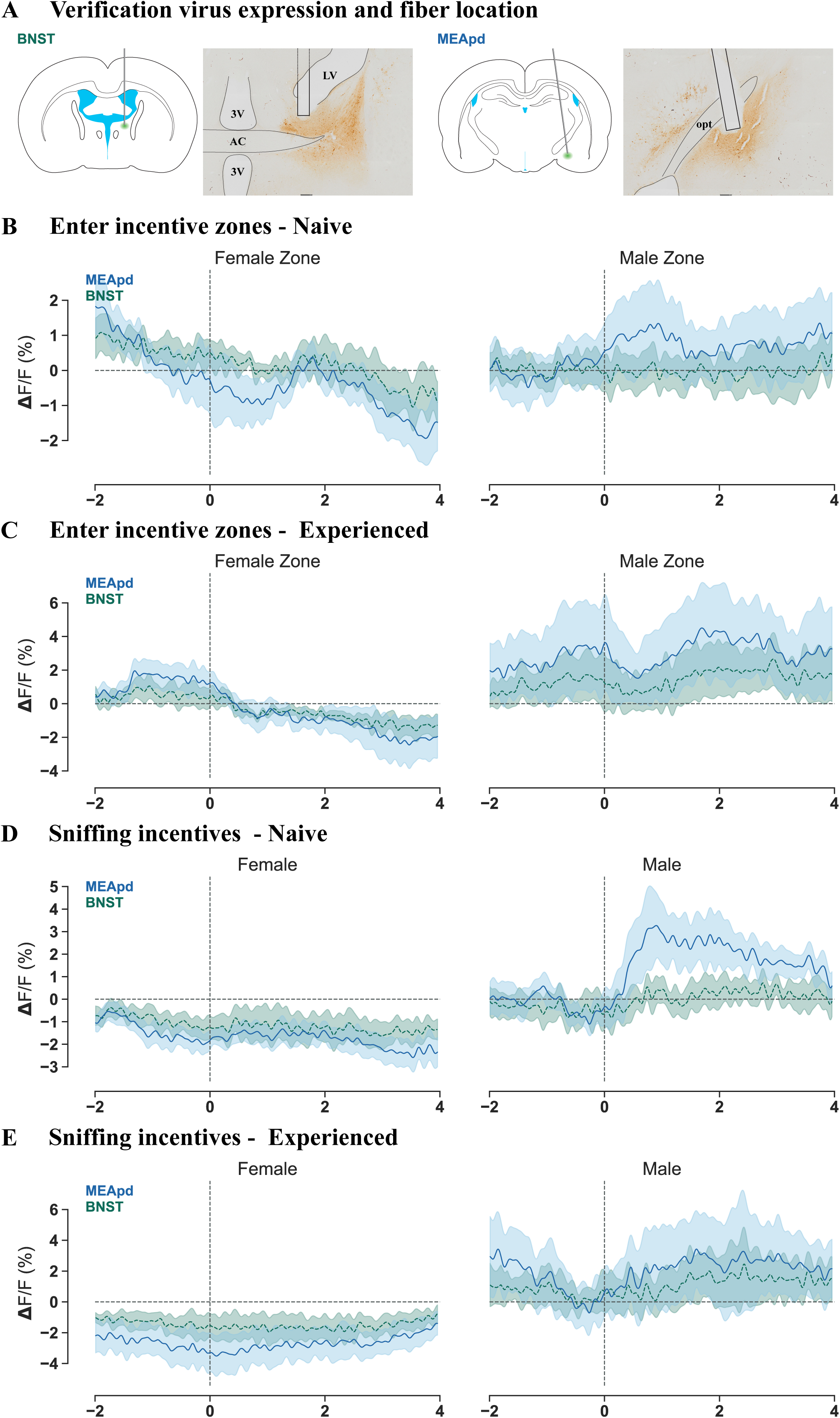
Neural dynamics in MeApd and BNST during incentive motivation test. A) Schematic representation of surgical details and histologically determined fiber locations and AAV vector expression in the BNST (left) and MeApd (right), B) ΔF/F aligned to the entry into the female (left) and male (right) zone in sexually naïve state, C) ΔF/F aligned to the entry into the female (left) and male (right) zone in sexually experienced state, D) ΔF/F aligned to sniffing the wire mesh in front of the female (left) and male (right) in sexually naïve state, E) ΔF/F aligned to the sniffing the wire mesh in front of the female (left) and male (right) in sexually experienced state. The lines in the ΔF/F figures represent the mean and shaded areas the SEM of GCaMP6s signaling in posterior dorsal part of the medial amygdala (MeApd, blue) and bed nucleus of stria terminalis (BNST, green). ac = anterior commissure, LV = lateral ventricle, 3V = 3^rd^ ventricle, opt = optic tract.

### MeApd and BNST activity during social and sexual incentive investigation

In the sexual incentive motivation test, males spent more time in the incentive zone adjacent to the receptive female than in the zone adjacent to the male stimulus, both before and after acquisition of sexual experience (Fig. S2A,F(3,18) = 11.707, p < 0.001; naïve: p = 0.001, experienced: p < 0.001). Preference scores similarly indicated a consistent behavioral preference for the receptive female in sexually naïve and experienced animals (Fig. S2B, naïve: t = 7.50, p < 0.01; experienced: t = 3.16, p = 0.02), with no difference between sexual-experience states (t = −0.18, p = 0.86).

Despite this behavioral preference, entry into the female- or male-associated incentive zones did not evoke reliable event-aligned changes in MeApd or BNST activity (Fig. 1B/C). AUC did not differ across zone-entry conditions in the BNST (Fig. S2C-F, F(7,42) = 1.022, p = 0.430), or MeApd, (Fig. S2C-F, F(7,42) = 1.079, p = 0.394). Similarly, sniffing through the wire mesh in front of the stimulus animals did not produce reliable AUC changes in either region (Fig. 1D-E, Fig. S2G-J, BNST: F(7,42) = 1.305, p = 0.272; MeApd: F(7,42) = 1.961, p = 0.084). Thus, indirect investigation across a wire barrier did not reveal robust region-specific activity, suggesting that direct social interaction may be required to detect stronger MeApd and BNST responses.

### Conspecific introduction recruits MeApd and BNST differently across social and sexual contexts

We next examined activity following introduction of a receptive female during copulation tests and an intact male during social interaction tests. Introduction of a conspecific increased MeApd activity across all conditions (Fig. 2A–C). MeApd AUC differed across condition and epoch (Fig, 2D-F, F(5,30)=18.251, p < 0.001), with post-event activity exceeding pre-event activity following introduction of a receptive female in sexually naïve males (adjusted p = 0.039), introduction of a receptive female in sexually experienced males (p = 0.001), and introduction of a male conspecific (p < 0.001). The post-introduction MeApd response to a male conspecific was larger than the response to a receptive female in both naïve and experienced conditions (both p < 0.001). Likewise, peak response amplitude in the MeApd also differed across social and sexual conditions (Fig. 2G, *F*(2,20) = 6.247, p = 0.008). Introduction of a male conspecific produced larger MeApd responses than introduction of a receptive female in both sexually naïve (p = 0.013) and sexually experienced males (p = 0.026).

**Fig. 2.**
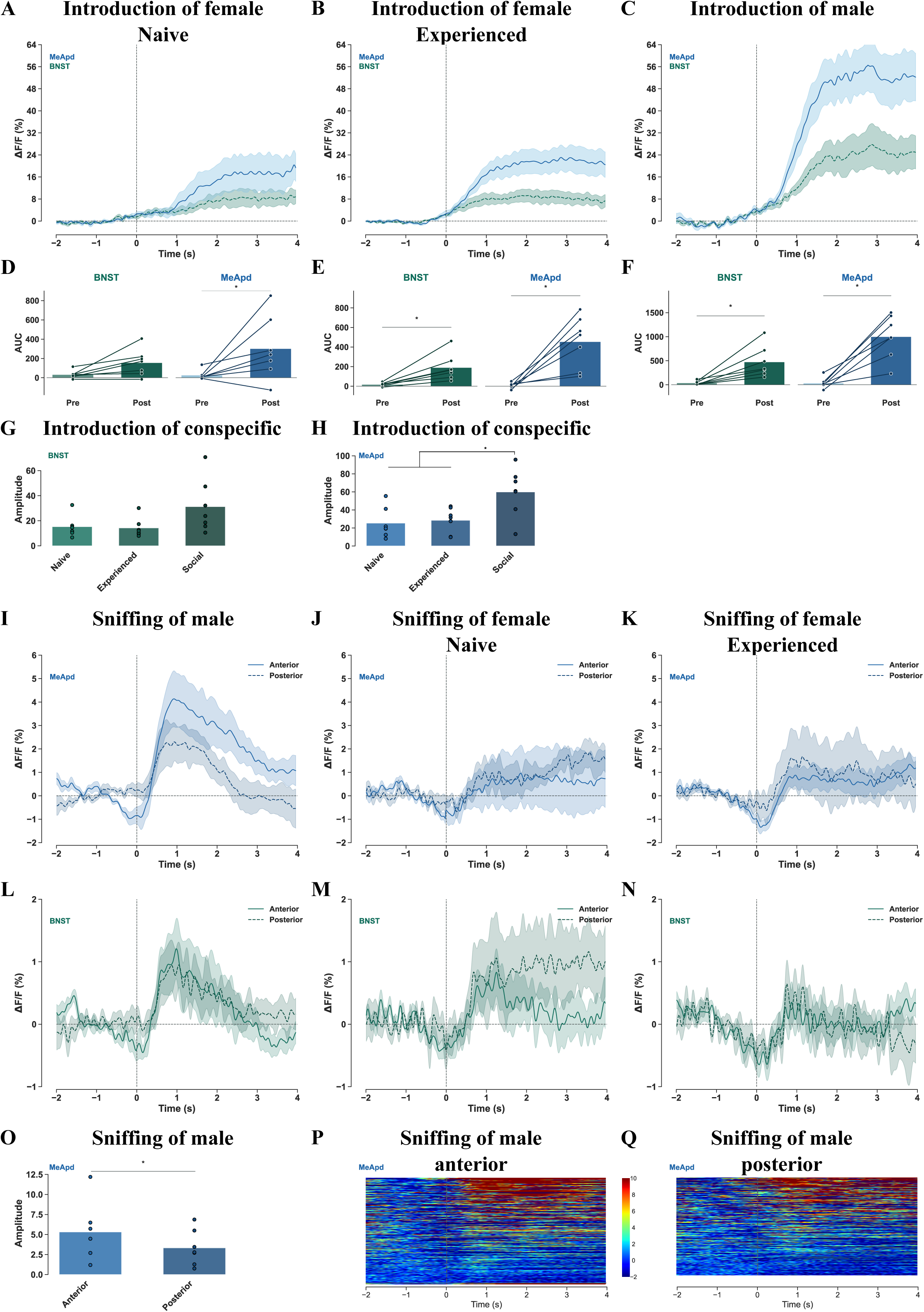
Neural dynamics in MeApd and BNST during olfactory investigation. A-C) ΔF/F aligned to the introduction of the receptive female (in sexually naïve or experienced state) or male conspecific, D-F) AUC from 2 seconds before (pre) and after (post) the introduction of the receptive female or male, G/H) amplitude of ΔF/F response to the introduction of conspecifics in the BNST and MeApd, I-N) ΔF/F aligned to the start of sniffing events of the anterior versus posterior parts of the body of the conspecific in the MeApd and BNST, O) amplitude of ΔF/F response upon sniffing the anterior versus posterior parts of the body of a male conspecific in the MeApd, P/Q) and heat maps of ΔF/F aligned to start of sniffing events. The lines in the ΔF/F figures represent the mean and shaded areas the SEM of GCaMP6s signaling in posterior dorsal part of the medial amygdala (MeApd, blue) and bed nucleus of stria terminalis (BNST, green). The dots and lines in the AUC plots represent data points from individual rats. * p<0.05 compared to by line connected group.

BNST responses showed a more experience-dependent profile (Fig. 2A–F). BNST AUC differed across condition and epoch (*F*(5,30) = 10.845, *P* < 0.001). Introduction of a receptive female did not produce a reliable pre-to post-event increase in sexually naïve males (p = 0.109), but did increase activity after sexual experience (p = 0.027). Introduction of a male conspecific also increased BNST activity (p < 0.001). Post-introduction BNST activity in the social condition was greater than in both the sexually naïve and sexually experienced female-introduction conditions (both p=0.002). The corresponding BNST amplitude comparison did not reach significance (Fig. 2H, *F*(2,20) = 3.296, *P* = 0.060).

### MeApd activity differentiates social investigation of anterior and posterior male body regions

Sniffing a receptive female in a sexual context did not elicit significant ΔF/F responses in either region, regardless of sexual experience (Fig. S3A-F). In contrast, MeApd activity increased during sniffing of a male conspecific in the social interaction test. MeApd AUC differed across social and sexual contexts (Fig. S3A-F, F(5,30) = 5.016, p = 0.002) with a significant pre-to post-sniffing increase only during social investigation (p < 0.001). Post-sniffing activity during social interaction was greater than during investigation of receptive females in sexually experienced animals (p = 0.014), whereas the comparison with sexually naïve animals was not significant (p = 0.240). BNST AUC did not differ reliably across these conditions (F(5,30) = 2.086, p = 0.095).

We next separated conspecific-directed sniffing according to the investigated body region. In the MeApd, investigation of the anterior head and face region elicited a larger response than investigation of the posterior anogenital and hind-body region (Fig. 2I-K, Fig. 2P/Q, and Fig. S3G-L). MeApd AUC differed across body region, condition, and epoch (F(11,66) = 2.950, p = 0.0039). During social interaction, anterior sniffing increased MeApd activity relative to baseline (p < 0.001), with a larger response than in both sexual conditions (social vs. naïve: p=0,014, social bs. experienced: p=0,013). The peak amplitudein a social context was also greater during anterior than posterior investigation (Fig. 2O, p = 0.046).

BNST activity did not show comparable body-region selectivity (Fig. 2L-N, and Fig. S3G-L). Although anterior investigation showed a trend toward increased AUC in the social condition (p = 0.051), the overall AUC model was not significant (F(11,66) = 1.342, p = 0.222). A modest amplitude difference was detected in sexually naïve animals (F(5,30) = 2.624, p = 0.044), in which posterior investigation elicited a larger response than anterior investigation (p = 0.011; Fig. 2P-Q). However, this effect was not accompanied by a corresponding AUC difference and was not consistently apparent in the event-aligned traces (Fig. S3G-L).

### MeApd and BNST activity increases during mounts and intromissions

We further investigated the functional involvement of the MeApd and BNST during copulatory behaviors, focusing on the impact of tactile stimulation intensity (mount, intromission, and ejaculation) and sexual experience on neural activity patterns. Mounts elicited increased calcium signals in both the MeApd and BNST in sexually naïve and sexually experienced males (Fig. 3A-E). In the BNST, AUC differed between pre- and post-mount epochs (F(3,18) = 5.374, p = 0.008), with significant increases in naïve (p = 0.010) and experienced males (p = 0.017). MeApd activity similarly increased during mounts (F(3,18) = 10.232, p < 0.001), in both naïve (p = 0.003) and experienced animals (p = 0.001). Event-aligned traces revealed biphasic responses during mounts, with a second ΔF/F increase occurring near mount termination and the onset of genital grooming (Fig. S4A-B). No reliable difference between sexually naïve and experienced conditions was detected for mount-associated activity in either region.

**Fig. 3.**
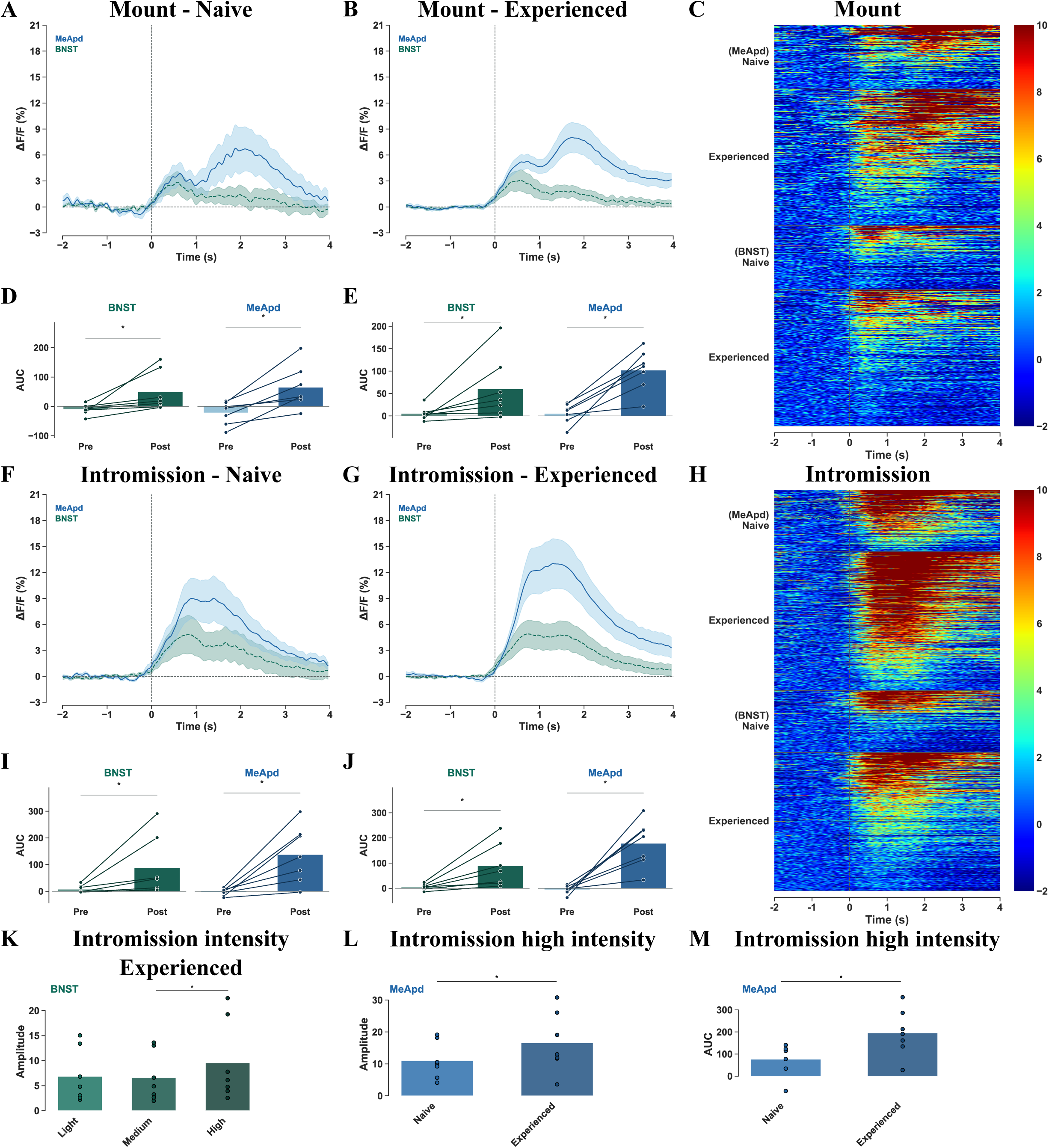
Neural dynamics in MeApd and BNST during tactile sexual stimuli with mounts and intromissions. A/B) ΔF/F aligned to the start of a mount in naïve or experienced state, C) heat maps of ΔF/F aligned to start of mount events, D/E) AUC before (pre) and after (post) the mount in naïve and experienced state, F/G) ΔF/F aligned to the start of a intromissions in naïve or experienced state, H) heat maps of ΔF/F aligned to start of intromission events, I/J) AUC before (pre) and after (post) the intromission in naïve and experienced state, K) amplitude of ΔF/F response to different intensity levels of an intromission in experienced state in the BNST, L) amplitude of ΔF/F response in naïve versus experienced rats upon the highest intensity intromission, M) AUC in response to the highest intensity intromission of naïve versus experienced rats. The lines in the ΔF/F figures represent the mean and shaded areas the SEM of GCaMPcs signaling in posterior dorsal part of the medial amygdala (MeApd, blue) and bed nucleus of stria terminalis (BNST, green). The dots and lines in the AUC plots represent data points from individual rats. AUC is calculated for 1 second before (pre) and 1 second after (post). * p<0.05 compared to by line connected group.

Intromissions also elicited increased activity in both regions (Fig. 3F-J). BNST AUC increased from pre-to post-intromission epochs in naïve and experienced males (F(3,18) = 5.478, p = 0.007; naïve, p = 0.013; experienced, p = 0.008). MeApd responses were similarly robust (F(3,18) = 14.586, p < 0.001; both naïve and experienced, p < 0.001). Additional traces showed that intromission-associated activity typically peaked near event termination and was often followed by a second peak (Fig. S4C-D).

When intromissions were categorized according to behaviorally observable intensity, all intensity categories were associated with increased AUC in both regions (Fig. S4E-L). In the BNST, the overall AUC model was significant with increased pre-post epochs in both naïve and experienced rats (F(11,66) = 4.463, p < 0.001, naïve: low intensity: p=0.010, middle intensity: p=0.013, high intensity: p=0.027; experienced: low intensity: p=0.006, middle intensity: p=0.006, high intensity: p<0.001), but AUC did not reliably differ between intensity categories. Peak amplitude in sexually experienced males, however, was greater during high-intensity than medium-intensity intromissions (F(5,30)=2.237, p=0.076, p = 0.038; Fig. 3K).

In the MeApd, increased pre-post epochs in AUC in both naïve and experienced rats were triggered (F(11,66) = 13.180, p < 0.001, naïve: low intensity: p<0.001, middle intensity: p<0.001, high intensity: p=0.008, experienced: low intensity: p<0.001, middle intensity: p<0.001, high intensity: p<0.001), but AUC did not reliably distinguish among intensity categories within sexual-experience states. However, responses to high-intensity intromissions differed between sexually naïve and experienced animals for both amplitude (F(5,30)=2.483, p=0.054, p = 0.009) and AUC (p < 0.001, Fig. 3L-M; Fig. S4K/M). Thus, BNST activity showed modest differentiation between intromission categories after sexual experience, whereas MeApd responses to high-intensity intromissions were modulated by sexual experience.

### Ejaculation is associated with the largest copulation-related responses

Ejaculation elicited robust increases in calcium activity in both regions (Fig. 4A-E). In the BNST, AUC increased from pre-to post-ejaculation epochs in naïve and experienced animals (F(3,18) = 6.178, p = 0.004; naïve, p = 0.013; experienced, p = 0.004). MeApd activity also increased reliably during ejaculation, (F(3,18) = 12.066, p < 0.001; both naïve and experienced, p < 0.001). Individual traces and heat maps demonstrated consistent ejaculation-aligned activity increases in both regions (Fig. 4A-C). Peak response amplitude during ejaculation did not differ between sexually naïve and sexually experienced animals in either the BNST (t = 0.392, p = 0.709, or MeApd, t = 1.040, p = 0.338, Fig, S5A-B).

**Fig. 4.**
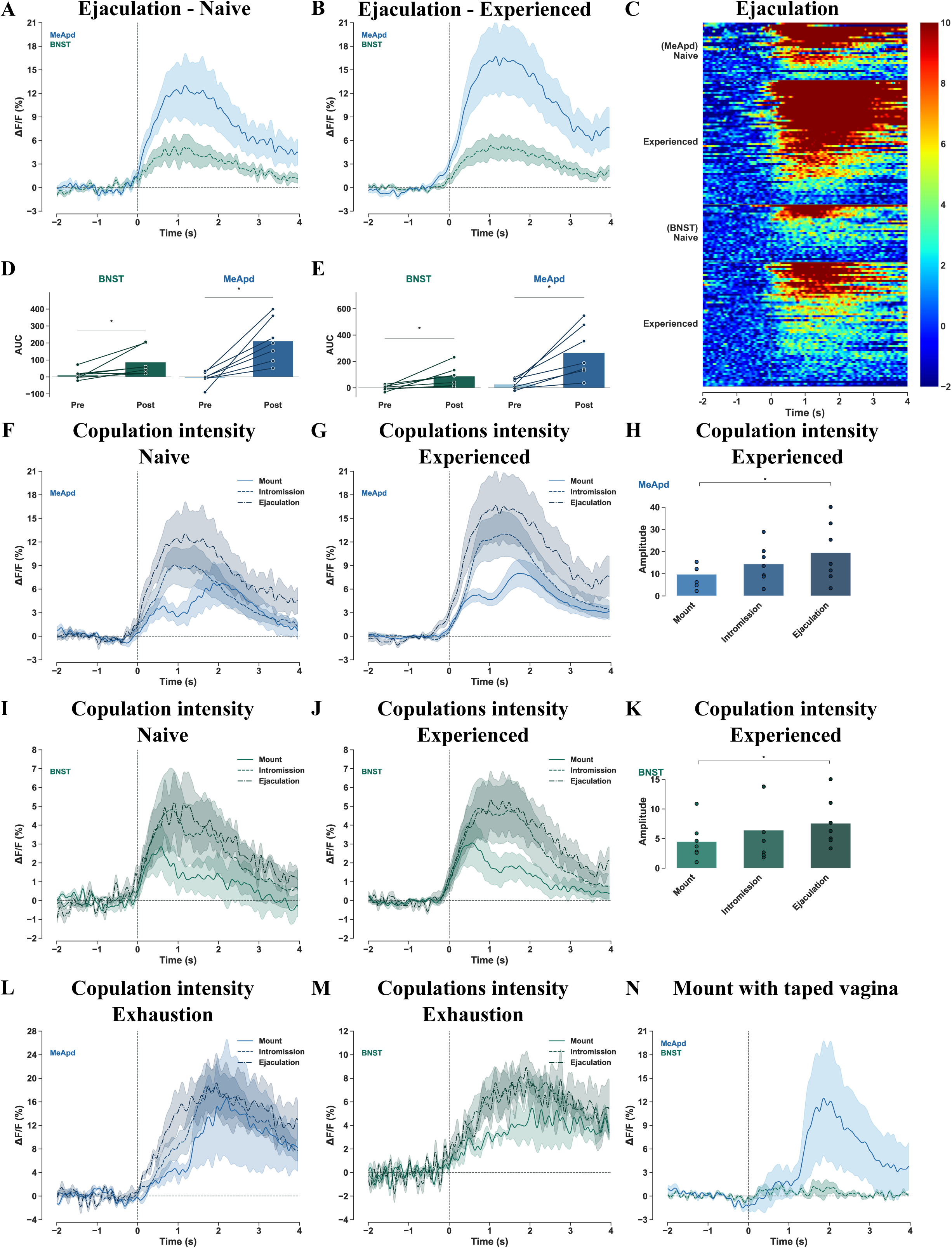
Neural dynamics in MeApd and BNST during tactile sexual stimuli with ejaculations. A/B) ΔF/F aligned to the start of ejaculations in naïve or experienced state, C) heat maps of ΔF/F aligned to start of ejaculations, D/E) AUC before (pre) and after (post) an ejaculation in naïve and experienced state, F/G) comparison of ΔF/F aligned to the start of mount, intromission and ejaculation in naïve or experienced rats in the MeApd, H) amplitude of ΔF/F in response to mounts, intromissions and ejaculations in MeApd, I/J) comparison of ΔF/F aligned to the start of mount, intromission and ejaculation in naïve or experienced rats in the BNST, K) amplitude of ΔF/F in response to mounts, intromissions and ejaculations in BNST, L/M) ΔF/F aligned to mounts, intromissions and ejaculations during the last ejaculatory series before exhaustion in MeApd and BNST, respectively, N) ΔF/F aligned to the start of mounts (lasting 1 second) with a receptive female with taped vagina. The lines in the ΔF/F figures represent the mean and shaded areas the SEM of GCaMP6s signaling in posterior dorsal part of the medial amygdala (MeApd, blue) and bed nucleus of stria terminalis (BNST, green). The dots and lines in the AUC plots represent data points from individual rats. AUC is calculated for 1 second before (pre) and 1 second after (post). * p<0.05 compared to by line connected group.

Across copulatory events, ejaculation was associated with the largest responses (Fig. 4F-K, Fig. S5C-H). In the MeApd, peak amplitude differed between event types and experience conditions (F(5,30) = 3.470, p = 0.014, Fig. 4H, Fig. S5E), with ejaculations producing larger responses than mounts in sexually experienced males (p = 0.009). MeApd AUC also differed across event types (F(11,66) = 11.858, p < 0.001, Fig. S5C-D), and post-event AUC was greater during ejaculation than mounting in both naïve (p = 0.002) and experienced animals (p < 0.001). In the BNST, peak amplitude differed across copulatory events (F(5,30) = 2.707, p = 0.039, Fig. 4K, Fig. S5H). After sexual experience, ejaculations elicited larger responses than mounts (p = 0.023; Fig. 4I-K). Although the BNST AUC model was significant (F(11,66) = 5.802, p < 0.001, Fig. S5F-G), post hoc comparisons did not identify reliable AUC differences between event types beyond the pre-to post-event increases within each condition.

### Analysis during sexual exhaustion

Since tactile sensations seem to matter, our next step was to investigate whether the neural responses of the MeApd and BNST would reduce over the course of multiple ejaculatory series, when rats reach exhaustion. Therefore, four rats were selected and allowed to copulate until they no longer showed any copulatory behavior for one hour^51,52^, and confirmed by the lack of copulatory behavior in the copulation test performed 24 hours later. During the final ejaculatory series preceding behavioral exhaustion, mounts were no longer associated with reliable increases in AUC in either region (BNST, p = 0.073; MeApd, p = 0.125; Fig. S5I/L). In contrast, intromissions and ejaculations continued to evoke increased activity in both regions. In the BNST (F(5,15) = 6.701, p = 0.002), AUC increased during intromissions (p = 0.003) and ejaculations (p = 0.002, Fig. 4M, Fig. S5J-K/M-N). In the MeApd (F(5,15)=8.114, p<0.001), AUC increased during intromissions (p = 0.008) and ejaculations (p < 0.001). Post-event MeApd AUC was greater for ejaculation than mounting (p = 0.014, Fig. 4L, Fig. S5O). Interestingly, the ΔF/F response upon ejaculation during this last series was similar as their 1^st^ ejaculation during the exhaustion test (data not shown)). Together this suggests that although the sensory information from tactile stimulations might reduce over the course of multiple ejaculation series, the BNST and MeApd remain robust in encoding copulatory behaviors. Interestingly, when re-exposed to a mate 24 hours later, one rat did show 1 mount and intromission. Although no conclusions can be drawn from this single observations, it is worth mentioning that the increase in ΔF/F was in line with what was previously observed (data not shown).

### Vaginal occlusion reduces copulation-associated activity

During copulation with vaginal occlusion, we did not observe enough extravaginial intromission-like patterns for analysis, but mounts were not associated with reliable event-aligned AUC increases in either region (Fig. 4N). For mounts, activity did not differ from baseline in the BNST (t = −1.344, p = 0.156), or MeApd (t = −0.646, p = 0.292). In contrast, the MeApd continued to show activity during the genital-grooming period that followed mounts, whereas BNST activity during this period was less apparent. These findings are consistent with a contribution of genital sensory feedback to MeApd and BNST activity during normal copulation.

### MeApd and BNST responses show largely overlapping timing

Finally, we compared response-onset latencies between regions to characterize the relationship between neural responses in the MeApd and BNST to explore potential functional interactions between these regions. No consistent regional difference in response onset was detected for conspecific introduction (region effect: F(1,30) = 0.979, p = 0.330; region × condition interaction: F(4,30) = 1.579, p = 0.205), mounts (region effect: F(1,21.532) = 0.012, p = 0.913; interaction: F(4,22.824) = 1.265, p = 0.313), intromissions (region effect: F(1,18) = 1.886, p = 0.187; interaction: F(2,18) = 2.468, p = 0.113), or sniffing events (region effect: F(1,30) = 0.968, p = 0.333; interaction: F(4,30) = 0.732, p = 0.577).

For ejaculation, the overall region effect did not reach significance (F(1,18) = 3.853, p = 0.065), but the region × experience interaction was significant (F(2,18) = 4.300, p = 0.030). In sexually experienced males, MeApd responses began earlier than BNST responses (p = 0.007). Thus, MeApd and BNST responses were largely temporally overlapping, although a modest experience-dependent difference in response onset was observed during ejaculation.

## Discussion

Using simultaneous dual-site fiber photometry in freely behaving male rats, we identified specialized yet temporally overlapping activity dynamics in the MeApd and BNST during social investigation and copulation. Three main findings emerged. First, MeApd activity was reliably associated with introduction and olfactory investigation of male conspecifics and differentiated anterior from posterior body investigation. Second, both MeApd and BNST showed robust activity aligned to mounts, intromissions, and ejaculations, with the largest responses occurring at ejaculation. Third, sexual experience preferentially altered BNST recruitment, including activity following introduction of a receptive female and differentiation across tactile copulatory events. Together, these findings support a model in which MeApd and BNST contribute distinct but overlapping activity patterns to olfactory and tactile cues during social and sexual behavior.

The MeApd showed strong responses following the introduction of both male and female conspecifics, but responses to males were larger. In addition, MeApd activity increased during direct investigation of male conspecifics, whereas sniffing of receptive females did not elicit similarly robust event-aligned responses. By contrast, the BNST showed activation during social sniffing, but did not particularly distinguish anterior from posterior investigation. BNST activity following introduction of a receptive female depended on sexual experience, with a reliable response emerging in sexually experienced but not sexually naïve males. This pattern suggests experience-dependent modulation of BNST population activity in response to sexually relevant social cues. The lack of response to a receptive female contradicts other studies that suggested that the MeApd and BNST respond stronger to olfactory cues from the opposite sex^53–56^. One explanation for this discrepancy could be the context of the studies. While others often test the neural responses to olfactory cues in an isolated setting, we recorded neural activity in a social and sexual behavioral test. Furthermore, while the other studies measure c-fos expression, which might represent a summation of several neural activations of non-distinct origin because of the time window that is necessary to induce c-fos to detectable levels, we recorded real-time activity from the neurons with fiber photometry. Under the present conditions, our findings suggest that MeApd activity was more consistently associated with male social investigation than with sexual investigation of receptive females.

Pheromonal signals are complex and do not just consist of a single cue identifying biological sex, but rather comprise mixtures of hormone-dependent volatile compounds and peptides^8,57–59^. Both sex and individual information can be relayed through these pheromones and are processed differently^58,59^. The stronger MeApd response to a male conspecific may therefore reflect the integrated salience of male social cues within the direct interaction context, rather than a simple preference for male over female odor. Likewise, the absence of robust responses during sniffing of receptive females does not imply that female olfactory signals are not represented in the MeApd. During copulation, females provide a complex and continuously changing combination of olfactory, tactile, and motivational cues, which may not be captured by alignment to individual sniffing events. In the BNST, on the other hand, recruitment is more strongly influenced by the learned or contextual significance of olfactory cues and is more likely to participate in downstream modulation of behavioral output rather than the initial sensory representation.

MeApd activity also differentiated investigation of anterior and posterior body regions of male conspecifics. This pattern is compatible with differential processing of body-region-specific social olfactory signals, because facial and anogenital regions can emit distinct odor blends and provide different forms of social information. This regional selectivity mirrors prior observations of stronger responses in the vomeranasal organs (VNO) to facial sampling than to anogenital sniffing^60^. Sensory neurons in the VNO access nonvolatile stimuli by direct contact with sources of pheromones^61,62^, and relay this information via the accessory olfactory bulb to the medial amygdala^10,63,64^. A parallel pattern of differential activation in the MeApd and the VNO is therefore unsurprising. By contrast, despite a similar projection from the VNO to the BNST ^9^, the BNST showed little activation during social sniffing and did not particularly distinguish anterior from posterior investigation. The MeApd and BNST are strongly interconnected and both participate in the broader social-behavior network, but their anatomical organization also supports partially distinct functions^11,14,65^. Our findings are therefore consistent with the possibility that MeApd activity more closely tracks features of direct social investigation, whereas BNST recruitment is more strongly influenced by the learned or contextual significance of olfactory cues and is more likely to participate in downstream modulation of behavioral output rather than the initial sensory representation.

When it comes to tactile cues, both MeApd and BNST showed robust activity aligned to discrete copulatory events. Mounts, intromissions, and ejaculations all elicited increased calcium signals, with ejaculation (most intense tactile stimulation during copulation) producing the largest responses. These findings extend previous immediate-early gene studies showing increased c-Fos expression in mating-related regions after sexual behavior, including stronger activation after intromissions and ejaculations than after mounts alone ^26^. Unlike these previous studies, which reflect neural activation across extended periods of sexual behavior, the present recordings show event-level activity dynamics associated with individual copulatory acts. We therefore showed that the act of an ejaculation itself activated the BNST and MeApd more strongly than mounts. In the MeApd, ejaculation elicited larger responses than mounting even in sexually naïve animals. In the BNST, differentiation among copulatory events was more apparent after sexual experience but then began to discriminate subtle intensity differences in intromissions. This graded response suggests that the MeApd and BNST are sensitive to the intensity of tactile feedback and that the MeApd encodes tactile salience early on, while BNST’s sensitivity to tactile intensity of copulation is enhanced by experience. The biphasic ΔF/F signals observed during mounts and intromissions, with the second peak coinciding with the end of the behavior and the onset of genital grooming, further highlight the temporal precision with which these regions process tactile information. This pattern may reflect the hedonic processing of genital contact and subsequent self-grooming, behaviors that are closely linked to the rewarding aspects of copulation^20,21^.

The vaginal-occlusion experiment provides additional support for a contribution of genital sensory feedback to MeApd and BNST activity during copulation. When vaginal penetration was prevented, mounts no longer elicited reliable event-aligned responses in either region, while genital grooming did. This result is consistent with the interpretation that activity during normal copulation is not explained solely by the gross motor pattern of mounting or intromission. Previous work has demonstrated that disrupted tactile feedback impairs sexual motivation and performance in both males and females^19,66–68^. The occlusion findings also support the view that mounts are not simply failed intromissions. Mounts normally involve bodily contact and may provide sensory feedback even in the absence of vaginal penetration (discussed before in^41^. However, when vaginal occlusion altered the sensory consequences of mounting, reliable event-aligned responses were no longer detected in either region.

The sexual exhaustion paradigm provided further insights into the adaptive nature of neural responses in the MeApd and BNST. In the final ejaculatory series before exhaustion, when one would expect the tactile sensation to become desensitized, neural responses to mounts indeed diminished, while responses to intromissions and ejaculations persisted. This suggests that sensory fatigue may reduce the salience of less intense tactile stimuli, such as mounts, while high-intensity stimuli like ejaculations remain robustly encoded, despite overall motivational decline^42^.

Sexual experience emerged as a critical factor in modulating the neural activity of mainly the BNST. In naïve rats, the BNST showed minimal response to the introduction of a receptive female, whereas a significant response emerged after sexual experience. BNST activity also showed greater differentiation between high-and medium-intensity intromissions, and between ejaculation and mounting, after sexual experience. These findings suggest that repeated copulatory experience changes the recruitment of BNST populations by sexually relevant cues and copulatory events, which may play a role in fine-tuning the temporal and motor aspects of copulatory behavior. This aligns with lesion and DREADD studies demonstrating that BNST inhibition prolongs ejaculation latency and increases the inter-intromission and time-out intervals preceding ejaculation^37–39^. This supports our previous suggestion that the BNST is involved in coordinating the pacing and efficiency of copulation, and the motivational aspects of re-engaging after copulatory pauses^39^. In comparison, MeApd activity was robust across sexually naïve and experienced conditions. This relative stability is consistent with the established role of the medial amygdala in integrating sensory information relevant to male sexual behavior^36^, and is consistent with evidence that sexual experience does not necessarily alter Arc expression in the MeA after mating ^23^. Nevertheless, MeApd responses to high-intensity intromissions differed between sexually naïve and experienced animals, indicating that MeApd activity is not fully independent of experience.

Despite their anatomical connectivity and functional interplay ^11,14,65^, and therefore the hypothesis that one region might influence the other, our data suggest that the MeApd and BNST encode social and sexual behaviors largely independently. Both regions exhibited simultaneous activation for most behaviors, with no consistent temporal delay between them. This suggests there is not an order of up or downstream signaling between these regions in the context of our study. The earlier MeApd response during ejaculation in sexually experienced animals may reflect more rapid recruitment of MeApd populations, but this interpretation remains tentative. Instead of confirming an interaction regulation of the brain region, however, this may just as well reflect the direct role of the MeApd in sensory integration ^36^, while the BNST’s slower response could be attributed to its modulatory role in temporal patterning of copulation, and the motivational aspects of re-engaging after copulatory pauses. ^39^. This functional independence challenges the view that the BNST merely relays outputs from the MeApd, instead supporting a model where both regions process distinct aspects of social and sexual stimuli in parallel.

While this study provides novel insights into the neural dynamics of the MeApd and BNST, several limitations must be acknowledged. Fiber photometry, though powerful, measures population-level calcium dynamics, which does not capture the activity of individual neurons or genetically defined cell populations. Moreover, variability in virus expression prevented direct comparisons of absolute fluorescence amplitude between the MeApd and BNST. This limited our ability to assess the relative contributions of these regions to specific behaviors, and only allowed us to look at the onset of neural responses. Future studies could address these limitations by employing cell-type-specific recordings and to target distinct neuronal populations (although proven more difficult in rats).

## Conclusion

Simultaneous recordings from the MeApd and BNST reveal specialized yet temporally overlapping activity dynamics during social investigation and copulation in male rats. MeApd activity was particularly associated with olfactory cues from investigation of male conspecifics and differentiated anterior from posterior social investigation. In contrast, BNST recruitment to receptive female olfactory cues and differentiation across tactile intensities of copulatory events were more strongly shaped by sexual experience. Both regions displayed robust activity aligned to copulatory events, with the largest responses occurring at ejaculation. These findings provide a framework for investigating how extended-amygdala circuits integrate social cues, genital sensory feedback, and experience during reproductive behavior.

## Acknowledgements

Financial support was received from a local AKM fund from UiT The Arctic University of Norway and a FRIPRO Young researcher grant from the Norwegian Research Council (#251320). We are grateful to Geed Assadi for her contributions with scoring the behavior, John Olav Folstad for analyzing the brains, and Sebastian Mai and Vi Tran for their help with determining the best method to process the neural signals. Furthermore, we thank Carina Sørensen, Izabela Szkotak, Lorenzo Ragazzi, Hallvard Haugen, Remi Osnes and Amalie Hofmeyer for the excellent care of the experimental animals, and Truls Traasdahl for the construction of the behavioral test boxes and the brilliant solution for our fiber photometry cables. Finally, we thank the Advanced Microscopy Core Facility of UiT The Arctic University of Tromsø for access to their equipment for the brain analyses.

## Author contributions

PTH: Experimental design, methodology, data gathering, data curation, writing – review and editing

RH: Experimental design, methodology, data gathering, data curation, analysis, supervision, writing – review and editing, funding acquisition

EMSS: Experimental design, methodology, data gathering, data curation, Programming/software, analysis, supervision, writing – original draft, funding acquisition

