## Supplementary data for "Specialized and experience-dependent neural dynamics in the medial amygdala and bed nucleus of stria terminalis upon social and sexual olfactory and tactile cues"

**A** **Verification of virus expression and fiber location in BNST**

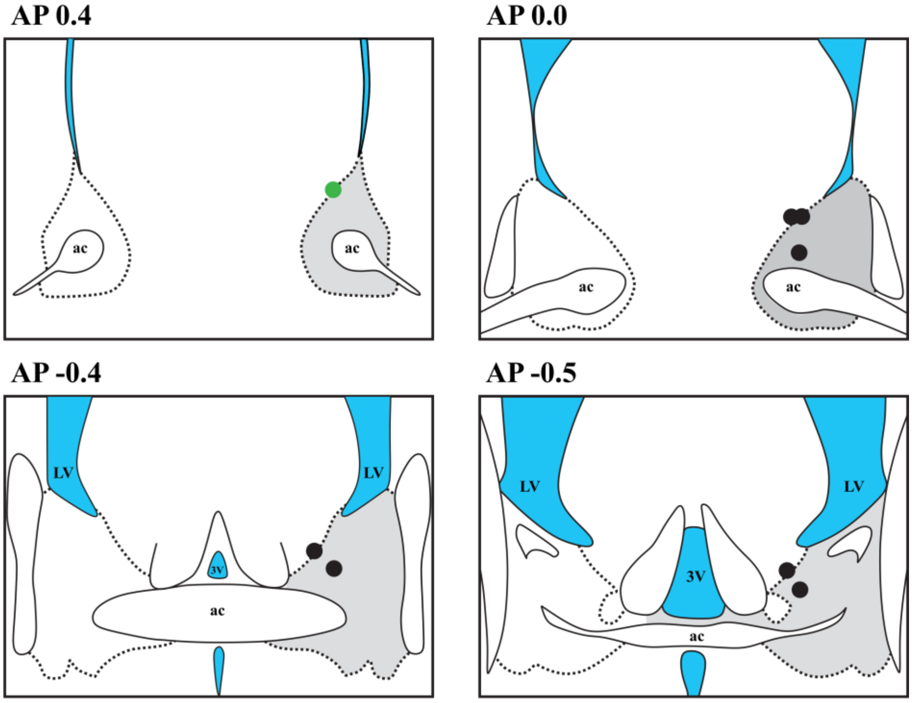


**B** **Verification of virus expression and fiber location in MeApd**

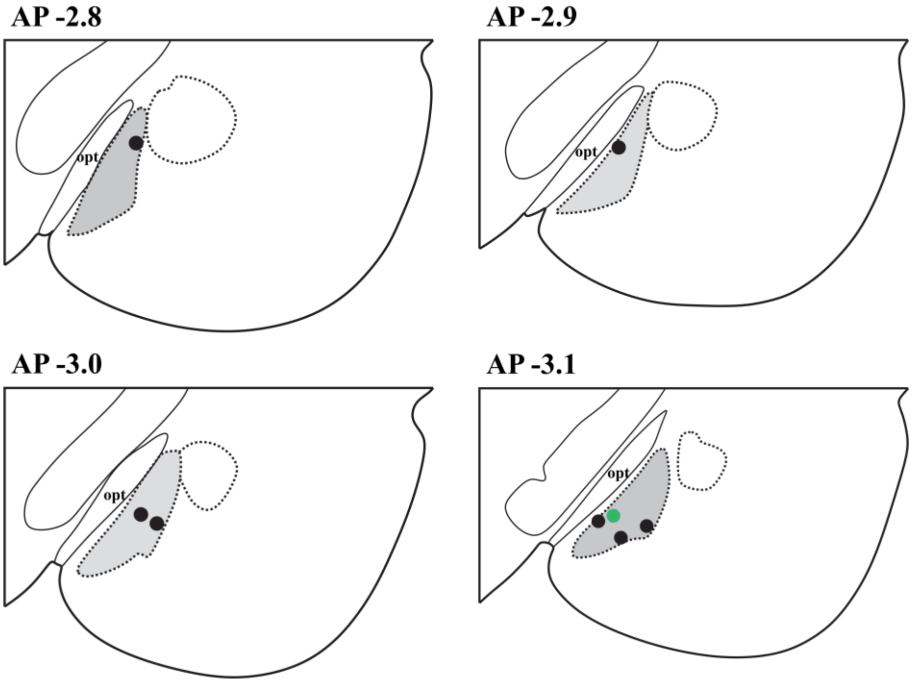


*Fig. S1. Schematic representation of surgical details and histologically determined fiber locations and AAV vector expression in the BNST (A) and MeApd (B). Individual rats with are marked with black (GCaMP6s) and green (eGFP) circles. ac = anterior commissure, LV = lateral ventricle, 3V = 3rd ventricle, opt = optic tract.*

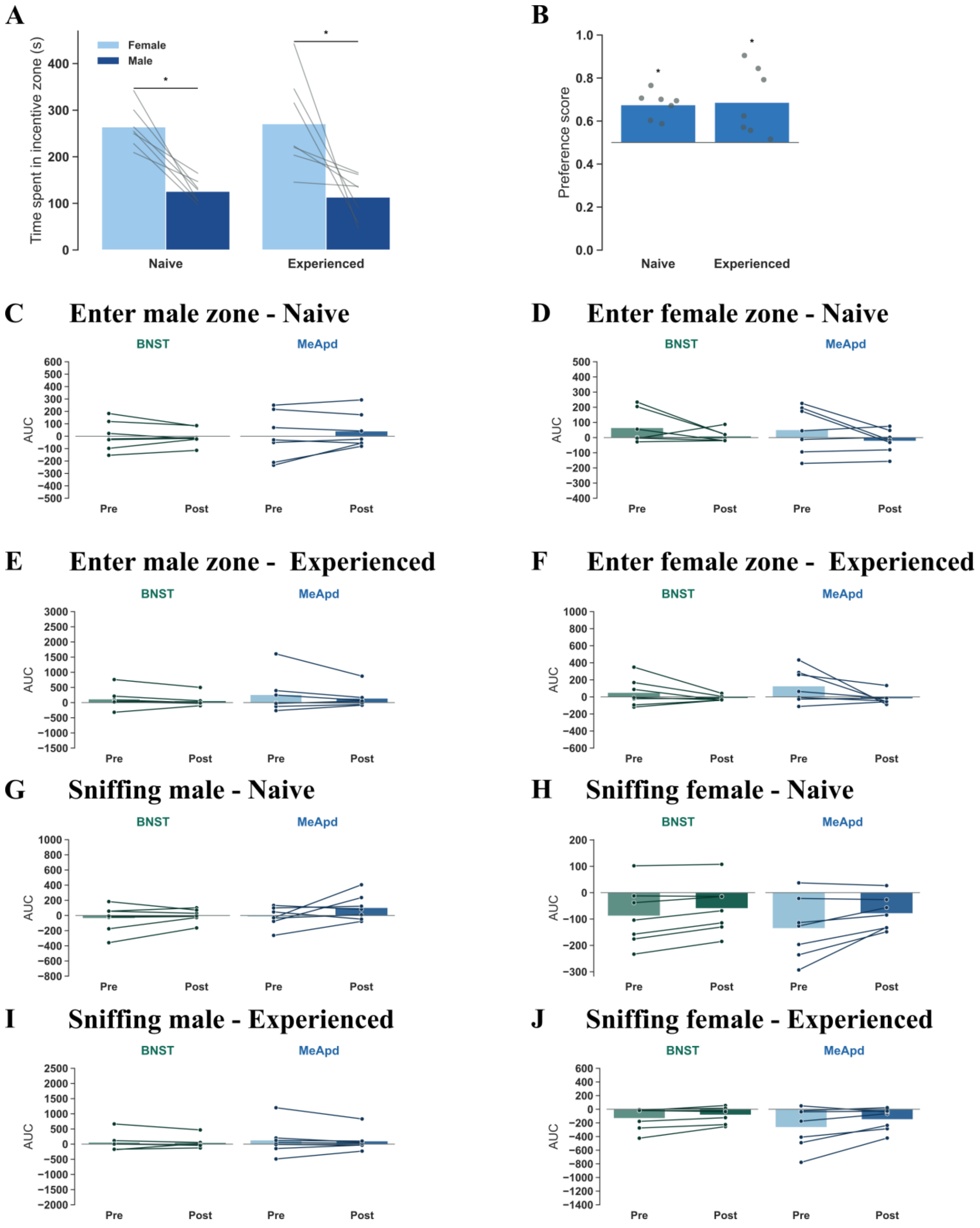


*Fig. S2. Sexual incentive motivation and neural dynamics in MeApd and BNST during incentive motivation test. A) Time spent in incentive zones with male or receptive female conspecific in sexually naïve or experienced state, B) preference score for receptive female in naïve or experienced state, C-F) AUC before (pre) and after (post) the entry into the male or female zone in sexually naïve or experienced state, G -J) AUC before (pre) and after (post) sniffing the grid of the male or female conspecific in sexually naïve or experienced state. The dots and lines in the AUC plots represent data points from individual rats. AUC is calculated for the 4 seconds before (pre) and after (post). * p<0.05 compared to by line connected group.*

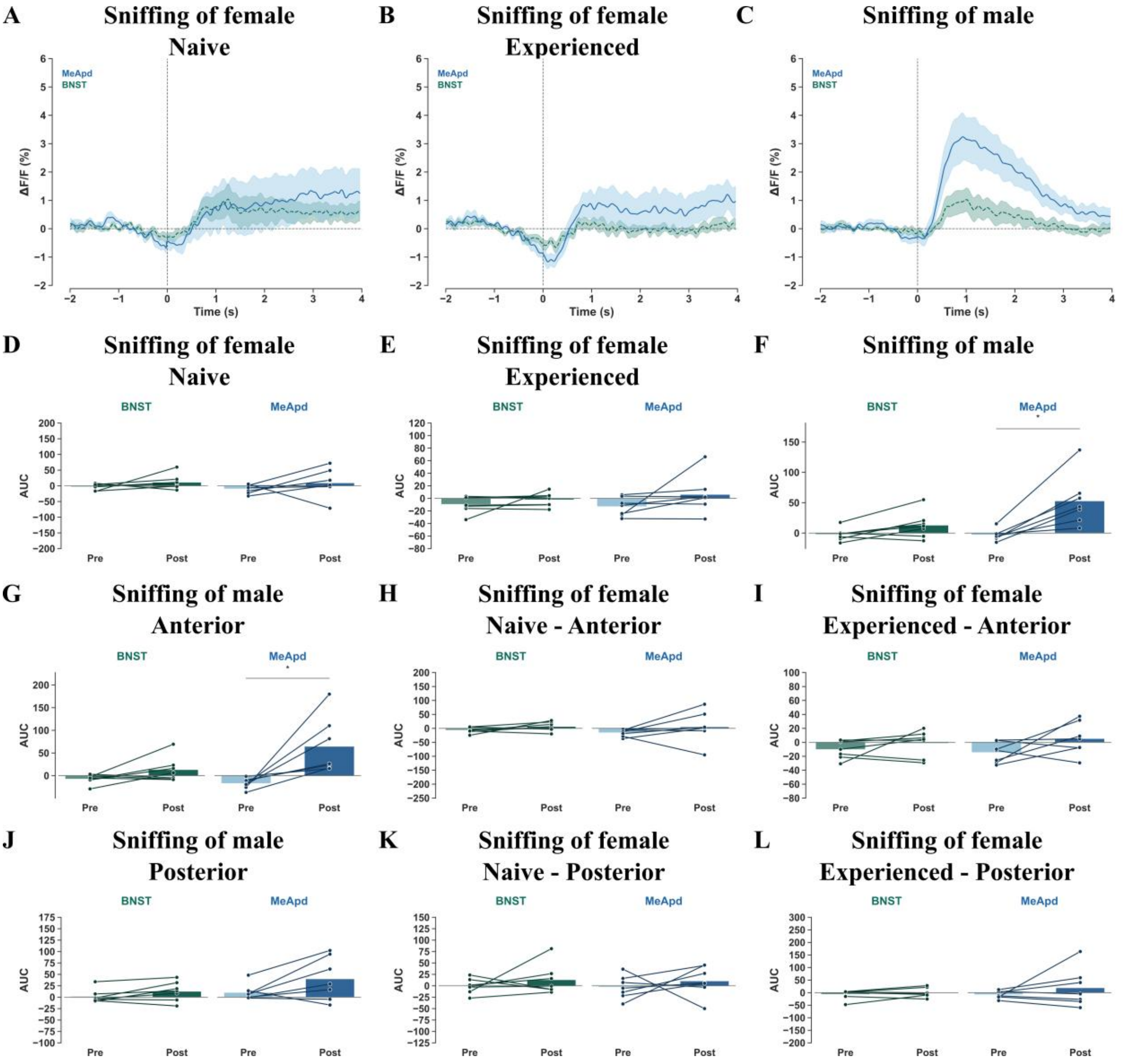


*Fig. S3. Neural dynamics in MeApd and BNST upon olfactory cues. A-C) ΔF/F aligned to the start of sniffing events in social, sexually naïve or sexually experienced context, D-F) AUC before (pre) and after (post) the sniffing the receptive female or male, G-L) AUC before (pre) and after (post) the sniffing the anterior or posterior part of the receptive female or male. The lines in the ΔF/F figures represent the mean and shaded areas the SEM of GCaMP6s signaling in posterior dorsal part of the medial amygdala (MeApd, blue) and bed nucleus of stria terminalis (BNST, green). The dots and lines in the AUC plots represent data points from individual rats. AUC is calculated for 1 second before (pre) and 2 seconds after (post), with the post corrected for proper comparison with pre. * p<0.05 compared to by line connected group.*

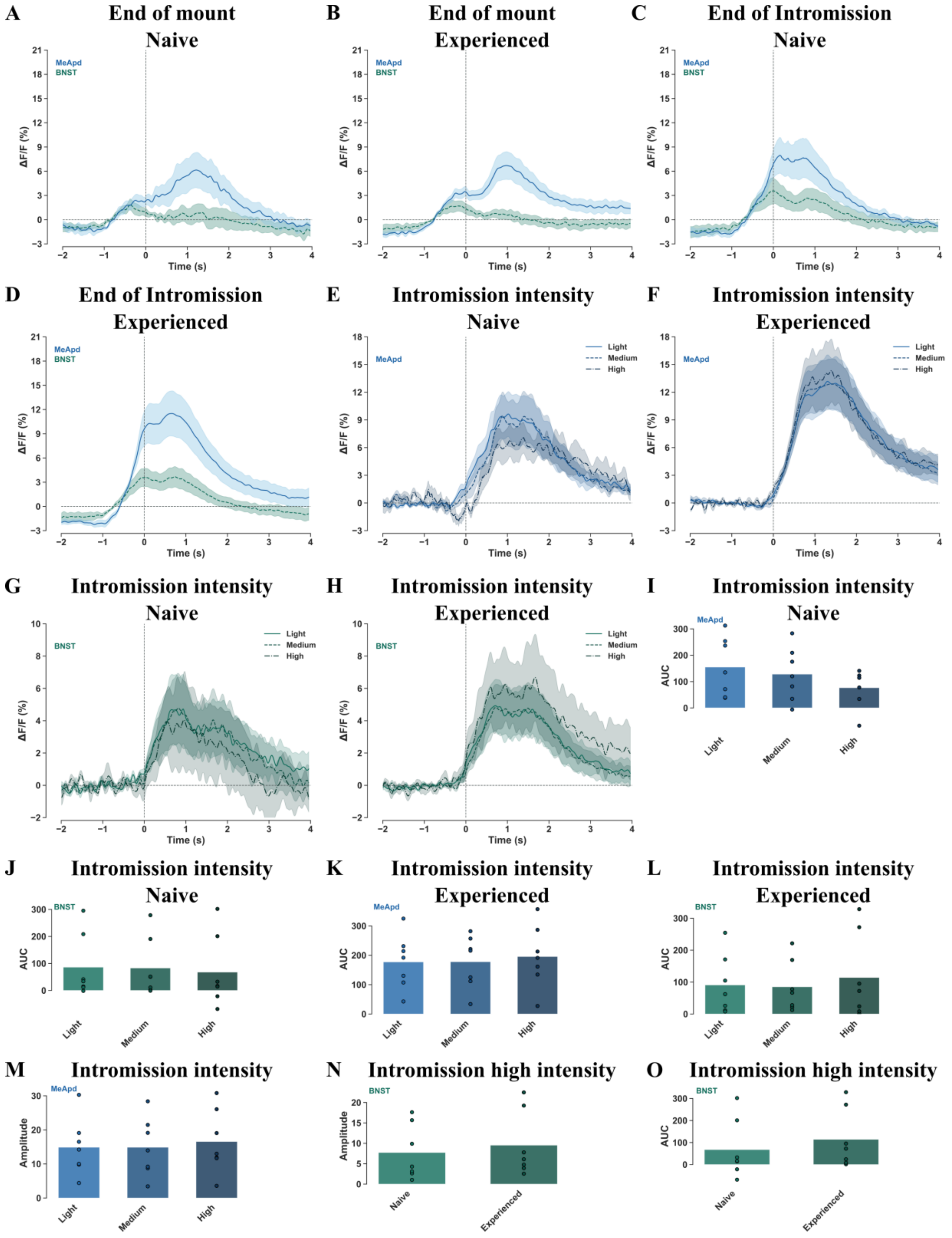


*Fig. S4. Neural dynamics in MeApd and BNST during tactile sexual stimuli. A/B) ΔF/F aligned to the end of a mount in naïve or experienced state, C/D) ΔF/F aligned to the end of an intromission in naïve or experienced state, E-H) ΔF/F aligned to light, medium and high intensity of intromissions in naïve and experienced states in the MeApd and BNST, I-L) AUC after light, medium and high intensity of intromissions in naïve and experienced states in the MeApd and BNST, M) amplitude of ΔF/F response to different intensity levels of an intromission in experienced state in the MeApd, N) amplitude of ΔF/F response upon highest intensity of*

*intromissions in naïve versus experienced rats in the BNST, O) AUC in response to the highest intensity intromission of naïve versus experienced rats. The lines in the ΔF/F figures represent the mean and shaded areas the SEM of GCaMP6s signaling in posterior dorsal part of the medial amygdala (MeApd, blue) and bed nucleus of stria terminalis (BNST, green). The dots and lines in the AUC plots represent data points from individual rats. AUC is calculated for 1 second before (pre) and 1 second after (post).* $p<0.05$ *compared to by line connected group.*

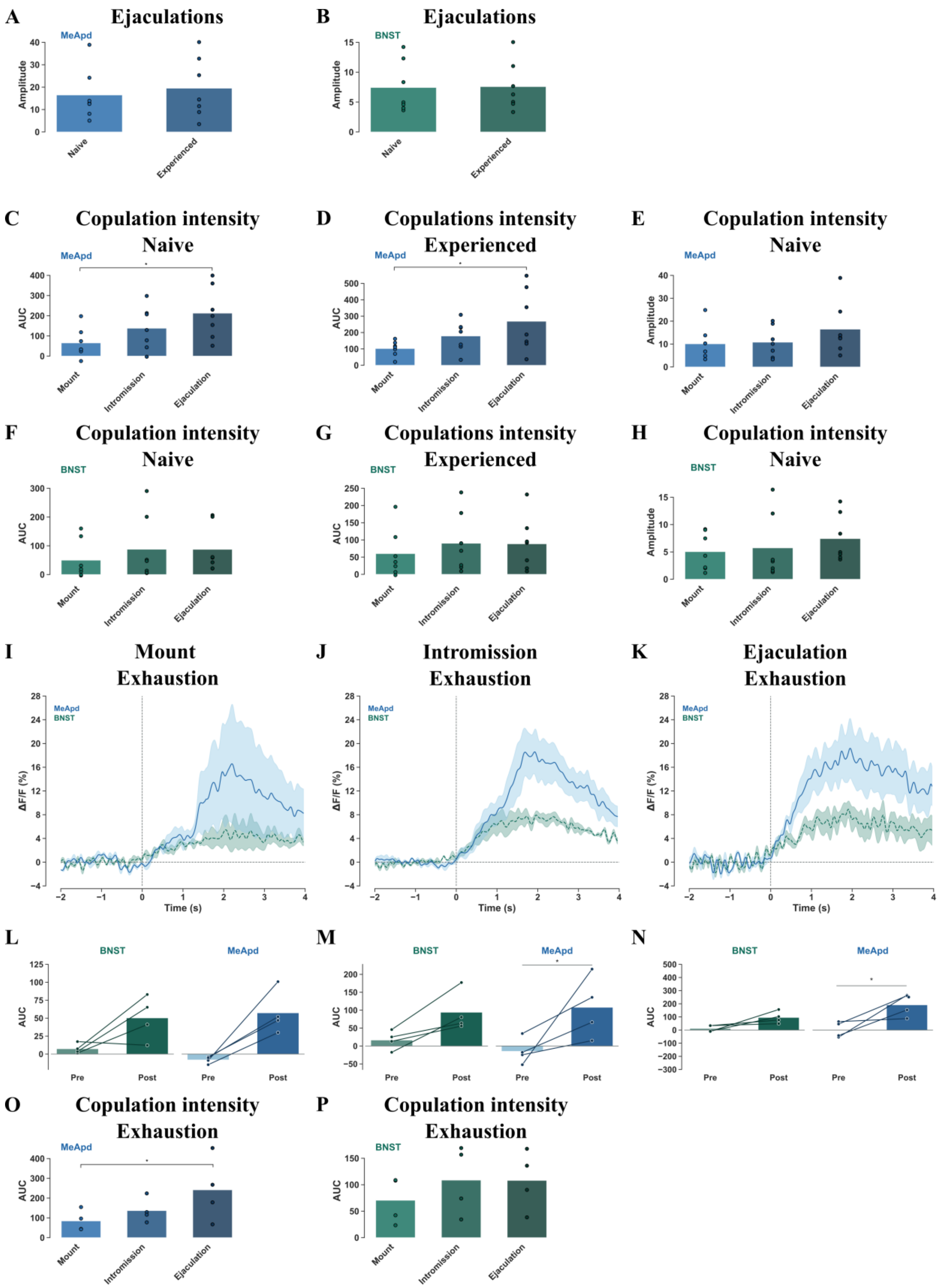


*Fig. S5. Neural dynamics in MeApd and BNST during ejaculations. A/B) amplitude of ΔF/F response to ejaculations in naïve or experienced state in the MeApd or BNST, C/D) AUC after mounts, intromissions and ejaculations in naïve or experienced states in the MeApd, E/) amplitude of ΔF/F response upon mounts, intromissions and ejaculations in naïve or experienced states in MeApd, F/G) AUC after mounts, intromissions and ejaculations in naïve or*

*experienced states in the BNST, H) amplitude of ΔF/F response upon mounts, intromissions and ejaculations in naïve or experienced states in BNST I) ΔF/F aligned to start of a mount, J) intromission, or K) ejaculation during the last ejaculatory series before sexual exhaustion, L-N) AUC before (pre) and after (post) mounts, intromissions and ejaculations during the last ejaculatory series before sexual exhaustion, O/P) AUC after mounts, intromissions and ejaculations during the last ejaculatory series before sexual exhaustion in the MeApd or BNST. The lines in the ΔF/F figures represent the mean and shaded areas the SEM of GCaMP6s signaling in posterior dorsal part of the medial amygdala (MeApd, blue) and bed nucleus of stria terminalis (BNST, green). The dots and lines in the AUC plots represent data points from individual rats. AUC is calculated for 1 second before (pre) and 1 second after (post). * $p<0.05$ compared to by line connected group.*
